# Fungal traits help to understand the decomposition of simple and complex plant litter

**DOI:** 10.1101/2022.12.06.519268

**Authors:** Eva F. Leifheit, Tessa Camenzind, Anika Lehmann, Diana R. Andrade-Linares, Max Fussan, Sophia Westhusen, Till M. Wineberger, Matthias C. Rillig

## Abstract

Litter decomposition is a key ecosystem process, responsible for the release and storage of nutrients and carbon. Soil fungi are one of the dominant drivers of organic matter decomposition, but fungal taxa differ substantially in their functional ability to decompose plant litter. We used a trait-based approach to better understand functional differences among saprotrophic soil fungi (originating from a natural grassland ecosystem) in decomposing leaf and wood litter. Decomposition strongly varied among phyla and isolates, with Ascomycota decomposing the most and Mucoromycota decomposing the least. In this study, the phylogeny of the fungi in our dataset, but also the ability of fungi to use more complex carbon were important predictors for decomposition. While some enzymes (e.g. laccase and cellulase) influenced decomposition, the majority of enzyme activities was not correlated with decomposition. Thus, we suggest using more directly assessed traits as predictors for decomposition, such as the ability to use carbon substrates, rather than a single enzyme activity, which could misrepresent the degradation potential of certain isolates. The findings of our study offer important new insights for the trait-based prediction of fungal litter decomposition in grassland soils.

## Introduction

Litter decomposition is a key ecosystem process, responsible for the release and storage of nutrients and carbon. Soil fungi are one of the dominant drivers of organic matter decomposition (Treseder and Lennon 2015). There is increasing evidence that fungal taxa differ in their functional ability to decompose organic matter, potentially contributing to large variability in this ecosystem process (Allison 2012; Crowther et al. 2014). Trait-based approaches have shown important functional differences among fungal lineages and have linked them to traits such as competitiveness, enzymatic capability, stress tolerance or growth rates (Lustenhouwer et al. 2020). These traits help to understand which strategies fungi develop to acquire their resources and how traits are linked to each other.

For instance, the production of certain enzymes can be limited while the fungus is investing in biomass gain. This trade-off has been shown for enzymes that come with a high metabolic cost, i.e. for laccase and cellobiohydrolase (Zheng et al. 2020). Generally, the relationship between litter decomposition and enzyme activity is not fully understood. In the past, it has been demonstrated that this relationship can be positive, neutral or negative (e.g. Allison and Vitousek 2004; Ge et al. 2017; Kang and Freeman 2009; Lustenhouwer et al. 2020). Most studies vary in their design, using either single or combined substrates and fungal species and sometimes measuring only a few single enzymes. Several studies have shown that the degradation of specific substrates requires the presence of specific functions, i.e. some substrates will only be decomposed if previous decomposition steps have released certain compounds, e.g. lignin is often decomposed in a later stage of decomposition (Algora Gallardo et al. 2021). Therefore, if one analyzes the decomposition of lignin in a short-term study that will only include early stages of decomposition, the mass loss of lignin will be unrelated or negatively related to the production of a certain enzyme. Similarly, if the *in vitro* decomposition of one single substrate by one fungal isolate is analyzed, the relationship might be negative, because the degradation of this substrate requires an enzyme repertoire that is provided by an entire fungal community when decomposition happens in the field.

Therefore, enzyme production only explains part of the variation in litter decomposition, e.g. Waring (2013) found that 35% of leaf decay rates could be predicted by enzyme activities. Sometimes, studies measured only a small number of enzymes or included only a limited number of organisms (e.g. Jiang et al. 2014, used three fungal species), thereby limiting their generalizability. In order to make this kind of generalization, we need more research on larger enzyme repertoires in fungal litter decomposition.

The functional ability to decompose litter further depends on the litter chemistry, such as the litter quality. Litter quality is usually subclassified in high quality litter, which is easily degradable, and low quality litter, which is difficult to degrade and more persistent (Castellano et al. 2015). Another important aspect of litter chemistry is the amount of water-soluble carbon in the litter (Allison and Vitousek 2004) and the identity and availability of particular carbon sources, which can influence the relative abundance of individual species and thus fungal community composition (Hanson et al. 2008). Litter decomposition further depends on the identity of the plant origin, the plant tissue (e.g. leaves or twigs), the number and the diversity of litter types present and also the nutrient status of litter and soil (Gartner and Cardon 2004; Grossman et al. 2020; Osono 2020; Porre et al. 2020; Yin and Koide 2019). For example, Yin & Koide (2019) describe in their study that the microbial community requires specific nutrients for the decomposition of more complex litter. Fungal decomposers have different strategies to deal with the challenges of varying nutrient and carbon (C) supply. Such options include a changing growth and respiration (C allocation), metabolism (e.g. enzyme excretion) and cell stoichiometry (Camenzind et al. 2021; Manzoni et al. 2021). Due to these functional differences among fungal taxa, the composition of microbial communities will be important to assess and forecast soil organic matter dynamics. Currently, only microbial biomass or fungal:bacterial ratios are incorporated into carbon cycling models. However, for the development of microbial-explicit models it will be important to understand the role of individual members and functional groups within fungal communities for litter decomposition and subsequent carbon release and storage (Hicks et al. 2022; Wan and Crowther 2022). So far, saprotrophic soil fungi have been studied with a focus on the decomposition of woody litter or dead wood in forest ecosystems (Bradford et al. 2014; Johnston et al. 2018; Osono 2020). Analyses of the ability of fungi to use different single carbon sources have also mostly been limited to fungi originated from forests or aquatic ecosystems (Algora Gallardo et al. 2021; Hanson et al. 2008; Sati and Bisht 2006).

Thus, with this study we aimed at better understanding functional differences among saprotrophic soil fungi originating from a grassland ecosystem, and their ability to decompose plant litter differing in complexity. Our set of fungi includes 31 isolates of the phyla Ascomycota and Basidiomycota, which are generally well known to be able to process various organic compounds including complex substrates, and Mucoromycota. For this last group this has been debated in the past, and a large study using fungi from forest ecosystems found that the ability of Mucoromycota to process complex litter is low compared with Asco- and Basidiomycota (Osono 2020). However, a recent study could show that these fungi, sometimes called ‘sugar fungi’, which generally grow well on glucose, have some families that also can process complex organic matter (Pawłowska et al. 2019).

We know from previous studies with our set of saprotrophic fungi that they have varying enzymatic capabilities, e.g. the 7 isolates of Mucoromycota included in this set are unable to produce laccase (Zheng et al. 2020). The 31 fungi also have varying aggregate formation capabilities, a key ecosystem process that is linked to many other processes, including stabilization and storage of organic matter (Lehmann et al. 2020). Moreover, these fungi have flexible stoichiometric reactions to differing nutrient supply, suggesting they can adapt to changing resource supply (Camenzind et al. 2021).

With this set of fungi, we performed three laboratory experiments in order to i) study their ability to decompose leaf and wood litter, ii) obtain further trait data on their ability to use single carbon substrates and iii) learn about their enzymatic capabilities. We additionally relate these trait data to an extensive dataset of existing chemical and biotic traits of the same set of fungi. We hypothesized that litter type will be an important factor for the weight loss of litter and that there will be strong differences among isolates, but also among phyla.

Especially for the Mucoromycota we expected low decomposition rates of woody litter. Similarly, we hypothesized that there will be important differences in carbon use and also enzyme activities among the fungal isolates; however, we did not expect each enzyme to correlate with decomposition as only two litter types and single isolates were used. Besides correlations of selected enzymes with decomposition, we rather expected to see correlations of sets of enzymes with the decomposition of complex C sources. For the use of carbon, we expected to see positive correlations between the use of the different carbon sources and litter decomposition, where carbon complexity is the main driver of those correlations.

## Material and Methods

### Fungal isolates

We used 31 fungal isolates previously characterized in detail (Andrade-Linares et al. 2016; Camenzind et al. 2022). Saprotrophic fungi were isolated from a grassland soil (Mallnow Lebus, Brandenburg, Germany, 52°27.778’ N, 14°29.349’ E) and cover the phyla Mucoromycota, Basidiomycota and Ascomycota (see Supporting Information, Tab. S1). Briefly, soil was washed and diluted in order to reduce spore abundance and to increase the quantity of living fungal hyphae attached to soil particles (Gams and Domsch 1967; Thorn et al. 1996). Subsequently, soil suspensions were spread on different media with the addition of antibiotics to prevent bacterial growth. Isolates were grown on potato-dextrose-agar (PDA) at room temperature thereafter (22°C).

### Experiment I: *In vitro* experiments on litter decomposition

We performed an *in vitro* study with soil and 31 saprotrophic fungal isolates decomposing two different types of litter (N=10 for each isolate), with 20 control plates (no fungus) per litter type, resulting in 660 experimental units. The stock of fungal cultures was grown on PDA and kept at 4 °C. For the experiment, we used 6 cm plates with water agar on top of which we added a 0.5 cm inoculum plug of the stock culture and placed a mesh bag (2×2 cm) next to the inoculum. The mesh bag contained 35 mg of dried litter, cut to 1 cm pieces. The litter types were linden wood (*Tilia cordata*) and leaves of a grass common at the field site (*Arrhenatherum elatius*). Finally, we added 10 grams of sterilized soil (autoclaved twice at 121°C for 20 min with a time delay of 24 hrs between events). The soil originated from the same field where the fungi were isolated. Soil properties were 0.11 % N, 1.45 % C, 19 mg kg^-1^ available P (Mehlich III extraction) and pH 6.79. The soil was wetted to 60 % field capacity. The plates were incubated in the dark for 10 weeks at room temperature. Upon harvest the soil and litter were dried and the litter weight loss was determined gravimetrically. We refer to litter decomposition as effect size calculated as

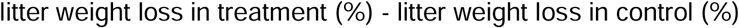

Hyphae were extracted from 4.0 g of soil (Jakobsen et al. 1992) and hyphal length (m g^-1^ soil) of all present hyphae in soil was measured according to (1999). We corrected the results for background concentration of hyphae from the control plates without fungi.

### Experiment II: *In vitro* experiments on carbon use

In a second experiment we tested the ability of the same set of fungi to utilize different carbon sources for growth (N=3; 30 fungal isolates (RLCS10 failed to grow on the medium type used); 540 experimental units). Growth media were designed following principles according to Liebig’s Law of the Minimum (Camenzind et al. 2020). All elements were provided in sufficient amounts to ensure C to be the only limiting element and to be able to observe effects exerted by C substrate manipulation. Element supply rates were determined based on the direct analysis of respective element contents in fungal tissues, avoiding osmotic stress by high salt additions (for details see Table S2 and Supporting Information S3). We prepared six different media using phytagel as gelling agent (2 g L^-1^) and final amounts of carbon compounds as follows: control (no C source), glucose 5 g L^-1^, cellobiose 4.75 g L^-1^, xylan (a hemicellulose) 4.4g L^-1^, cellulose 4.5 g L^-1^, and litter 4.42 g L^-1^. Litter represented a mixture of plant litter collected at the site of fungal isolation, was dried at 40°C and ground to a powder. C additions were standardized by molar amounts, resulting in comparable concentrations of C for each treatment.

The relative biomass gain of the fungi compared to the control treatment on each C substrate was our response variable. The use of a carbon source *j* for a fungal isolate *i* was calculated as follows, based on average values of all 3 replicates:

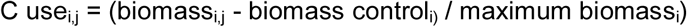

For this experiment we also calculated an indicator for the ability of fungal isolates to use more complex C compounds, hereafter referred to as “complex carbon use ability” (CCUA), which we defined as the weighted average of values according to the complexity of C compounds used:

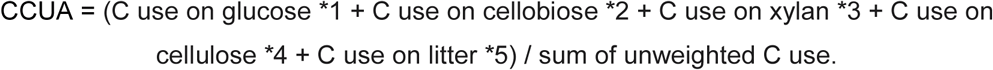

### Experiment III: In vitro experiments on enzyme activity

We determined the activity of a number of hydrolytic enzymes using the laboratory kit API ZYM™ (BioMerieux) for semi-quantitative analysis of selected enzymes produced by microorganisms. Fungi were grown on PDA and fungal material was collected from the margin of an actively growing fungus, the agar was removed and the fungal biomass homogenized with 2 ml of NaCl solution (0.85 %). Following the manufacturer’s instructions, 65 µl of the resulting suspension were added to each cupule of the kit and incubated at 37°C for 4.5 hours. The intensity of enzyme activity (nmol substrate hydrolyzed) was taken as response variable. We tested for activity of acid phosphatase, alkaline phosphatase, esterase, lipase, leucine arylamidase, phosphohydrolase, valine arylamidase, trypsin, α-galactosidase, α-chymotrypsin, α-glucosidase, α-mannosidase, β-galactosidase, β-glucoronidase, β-glucosidase and N-acetyl-β-glucosaminidase. As control, the test was run with only agar from the PDA medium.

### Trait collection

To complete our trait database, we selected further traits from previously published data using the same set of 31 fungi (see Tab. S1; Lehmann et al. 2020; Zheng et al. 2020). We selected traits that likely influence litter decomposition or can be linked with decomposition, either through biotic properties of the fungus (e.g. colony extension rate) or through chemical properties (e.g. enzyme activities).

**Table 1:** Traits used from the trait database.

| Trait | Description |
| --- | --- |
| Aggregate formation | Ability to form new soil aggregates (Lehmann et al. 2020) |
| Biomass density | Biomass density on potato dextrose agar ($\mu\text{g mm}^{-2}$) (Lehmann et al. 2020) |
| Colony extension rate | Colony growth rate on potato dextrose agar ($\mu\text{m h}^{-1}$) (Lehmann et al. 2020) |
| Competitive ability | Sum of wins against competitors in pairwise interactions (Soliveres et al. 2018) |
| Hydrophobicity | Hydrophobicity of fungal surface (Alcohol droplets test) (Lehmann et al. 2020) |
| Cellobiohydrolase, laccase, leucine aminopeptidase, acid phosphatase | Enzyme activity measured on fungal material growing on potato dextrose agar ($\text{U mg}^{-1}$ biomass) (Zheng et al. 2020) |

### Statistical analyses

We used the software R for all statistical analyses (version 4.1.3 R core team, 2021).

As a first step, we visualized the raw weight loss data using swarm plots and an estimation method that generates unpaired mean differences (treatment minus control) with bootstrapping (5000 iterations, package “dabestr”) (Ho 2022, Ho et al. 2019). This plot allowed us to see the magnitude and precision of the fungal taxa effect on litter decomposition compared to the control. The bootstrap estimates are by default bias- and skewness-corrected and produce 95% confidence intervals (CI), which, if situated above the zero line (line of no effect), can show that the treatment induced a detectable response.

As a second step, we visualized raw decomposition data (treatment - control) for individual isolates, and subsequently analyzed mean decomposition of each isolate (mean of 10 replicates) for correlations with carbon use (data from experiment II) and enzyme activities (experiment III) and traits of our trait collection using Spearman’s rank correlations (Spearman’s ρ). We further tested correlations of mean decomposition of each isolate for phylogenetic signals using phylogenetic generalized linear models (pgls function in package ‘caper’; Orme 2018).

Thirdly, we performed a principal component analysis to learn about links of decomposition with other variables of the whole trait dataset available for this set of fungi. In order to increase readability of the PCA we reduced the number of traits included in the analysis and thus selected those traits for the PCA that showed a significant relationship with decomposition in the Spearman or the pgls analysis.

## Results

### Litter decomposition

Across all isolates, weight loss of litter was 55% for leaf litter and 13.8% for wood litter, while in the control weight loss was 35.5% and 4.9%, respectively (See Fig. 1a and b). Weight loss across Ascomycota was on average 61.7% for leaf litter and 18.9% for wood litter; Basidiomycota caused an average weight loss of 64.5% in leaf and 17.5% in wood litter, and Mucoromycota had an average weight loss of 38.7% for leaf and 5.1% for wood litter. Both Asco- and Basidiomycota had CIs above the zero line for leaf and wood litter, i.e. they actively decomposed both litter types, whereas Mucoromycota had CIs overlapping with the zero line, showing a neutral response.

**Fig. 1.**
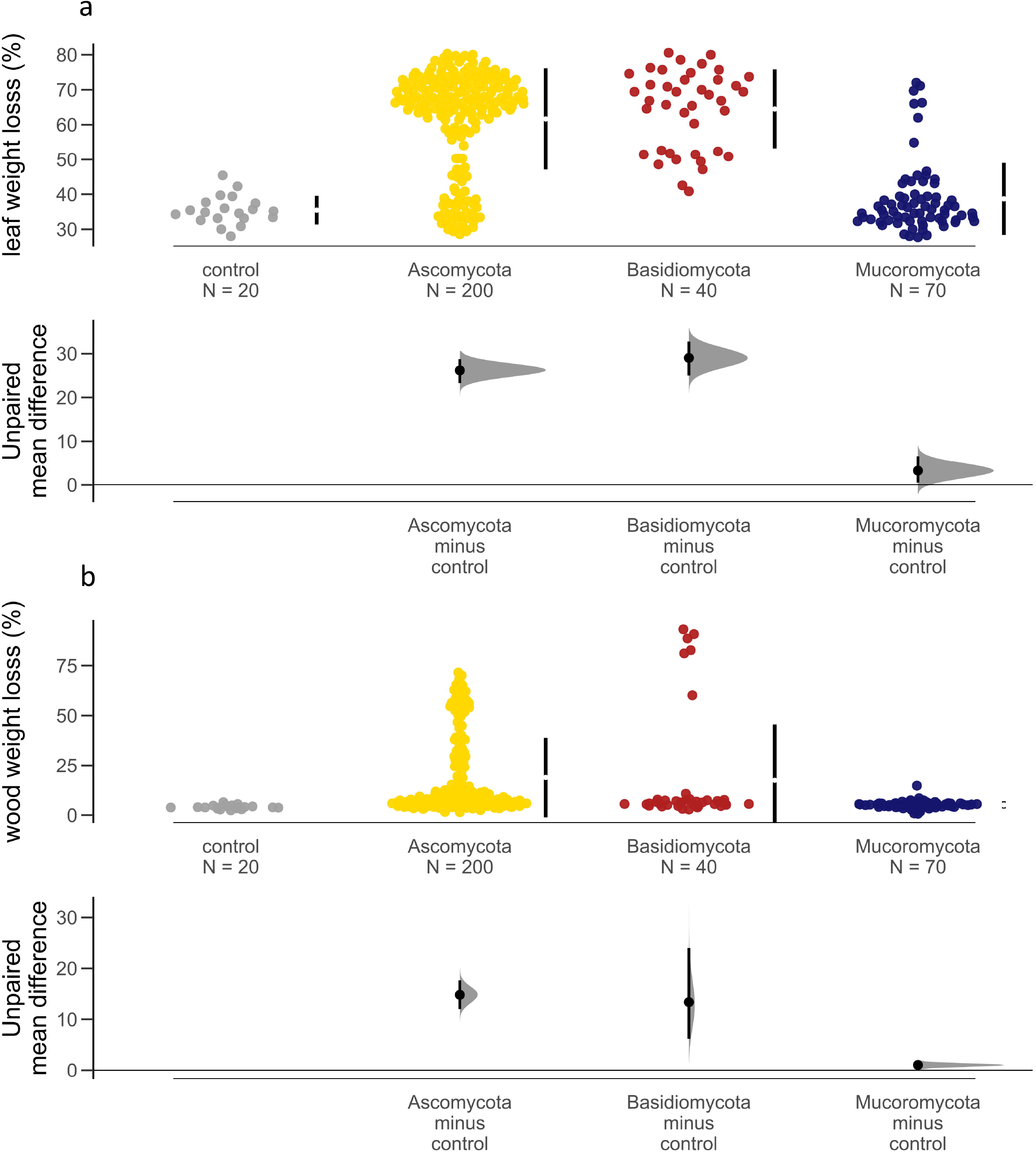

Decomposition (treatment - control) varied substantially among isolates (see Fig. 2a and b). For leaf litter there was also variation among the replicates of one isolate, for example for RLCS01 (*Mucor fragilis*), RLCS02 (*Mortierella elongata strain 2*) and RLCS04 (*Mortierella exigua*) there were some replicates that decomposed litter and some not at all. These strains are all within the Mucoromycota, which decomposed on average only slightly more than the control (plus 3.2%). One strain (RLCS09, *Trametes versicolor*) had considerable variation in decomposition of both leaf and wood litter. The leaf and wood litter in the control plates also had weight loss, which can be explained by degradation through fragmentation caused by wetting the experimental units and through handling the litter at harvest. Therefore, a few replicates showed negative results for decomposition.

**Fig. 2.**
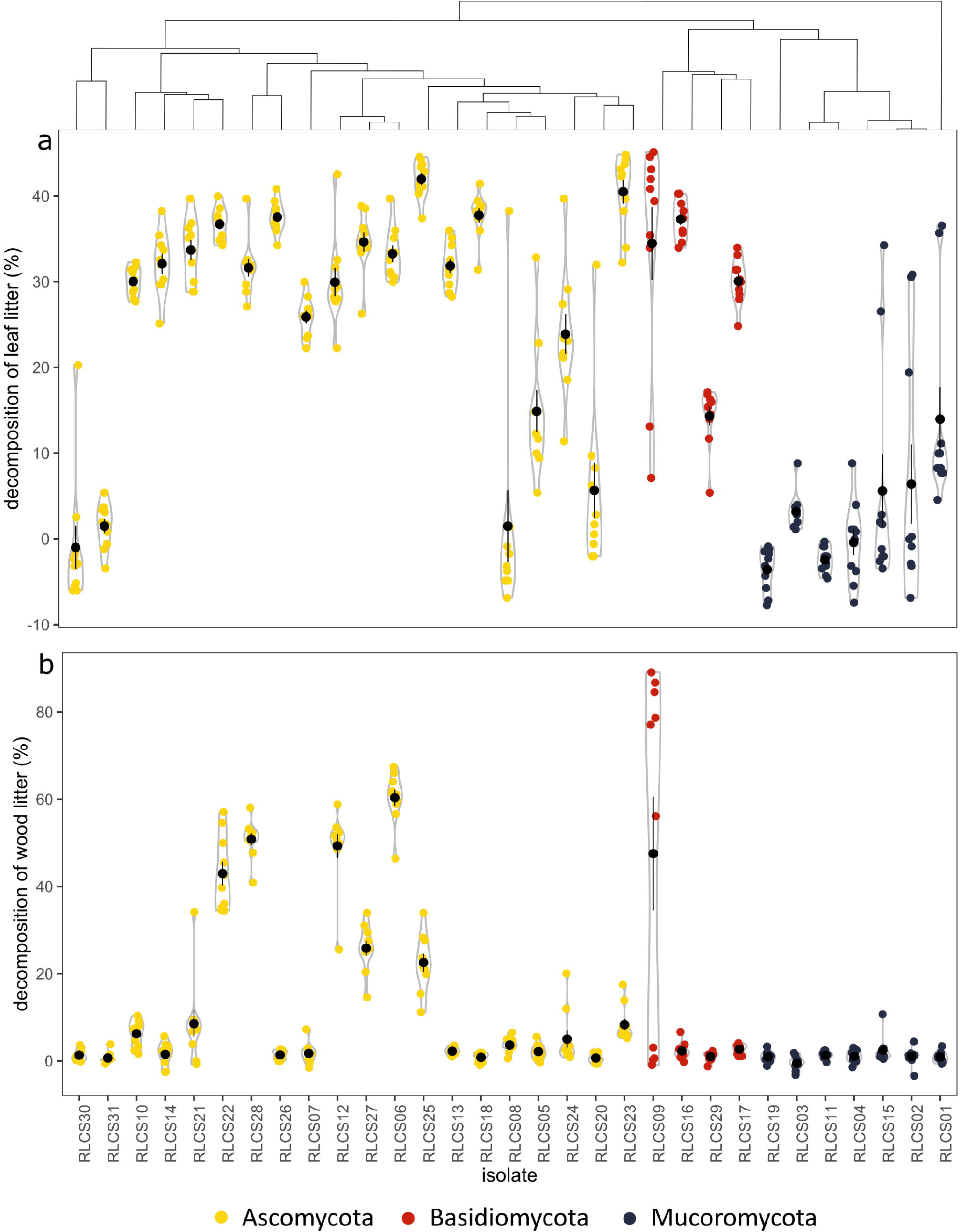

### Carbon use

The carbon sources were preferred in the order glucose > cellobiose > xylan > cellulose > ground litter (see Fig. 3a). Glucose was used by almost all fungi very well, while pure cellulose and ground litter were not used by the majority of the isolates. However, we also found differences between isolates, e.g. RLCS29 (*Macrolepiota excoriata*, Basidiomycota), RLCS12 (*Didymellaceae*, Ascomycota) and RLCS06 (*Chaetomium angustispirale*, Ascomycota) were able to use the more complex substrates comparatively well, while they could not use the glucose as much as the other isolates.

**Fig. 3.**
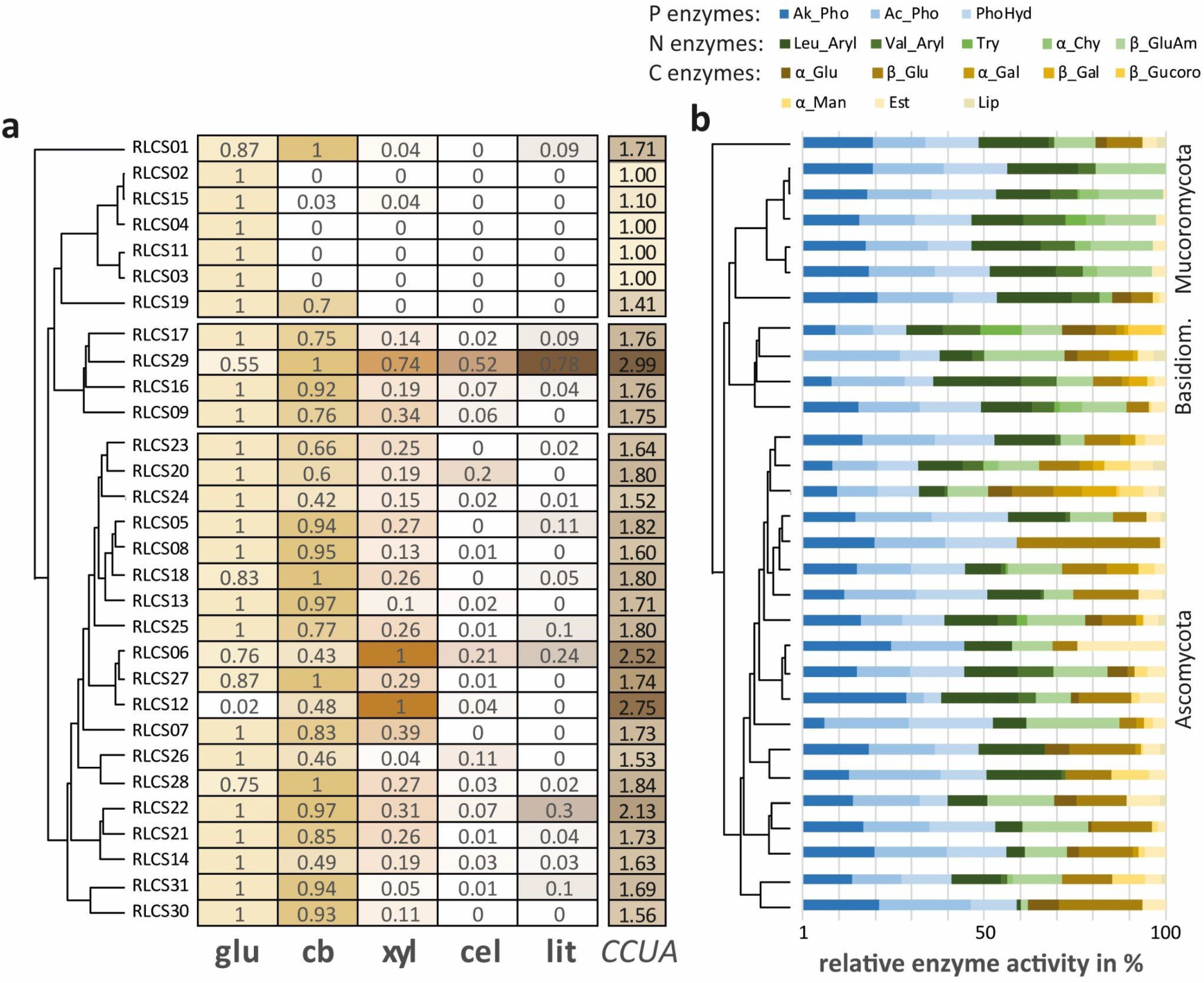

### Enzyme activities

A number of enzymes were produced by >70% of isolates, although in contrasting intensities. Among them were alkaline phosphatase, acid phosphatase, esterase, leucine arylamidase, valine arylamidase, phosphohydrolase, β-glucosidase, N-acetyl-β-glucosaminidase were very abundant (see Fig. 3b). Activity of lipase, trypsin, α-chymotrypsin, α-galactosidase, β-galactosidase and β-glucuronidase was detected in very few species (≤10 isolates).

The **Mucoromycota** showed a more uniform enzyme pattern compared to the other phyla. Their enzyme repertoire mainly consisted of acid and alkaline phosphatase, phosphohydrolase as well as leucine and valine arylamidase. Basidiomycota and Ascomycota had more diverse enzyme repertoires and large variations among their isolates. In **Basidiomycota**, for all isolates of our set, activities of the following enzymes were detected: alkaline phosphatase, leucine arylamidase, valine arylamidase, acid phosphatase, phosphohydrolase, β-glucosidase and N-acetyl-β-glucosaminidase. In **Ascomycota**, for most isolates of our set, activities of the following enzymes were detected: alkaline phosphatase, esterase, leucine arylamidase, acid phosphatase, phosphohydrolase, N-acetyl-β-glucosaminidase and β-glucosidase.

When we correlated all traits across our set of 31 fungi the use of hemicellulose (xylan) had the strongest relationship with the decomposition of leaf and wood litter (leaf: Spearman’s ρ = 0.54, p-value = 0.002; wood: Spearman’s ρ = 0.67, p-value = < 0.0001; Fig. 4). Leaf decomposition also had strong links with laccase activity and CCUA (laccase: Spearman’s ρ = 0.53, p-value = 0.002; CCUA: Spearman’s ρ = 0.51, p-value = 0.004). Cellulase activity and the ability to form new aggregates showed slightly less strong positive links with leaf decomposition. α-chymotrypsin showed a negative relationship with both litter types and had the second strongest link with wood decomposition (Spearman’s ρ = 0.56, p-value = 0.001). When we considered phylogenetic dependencies of the data in the pgls models we saw a different picture for leaf and wood decomposition. For leaf decomposition the activity of leucine arylamidase had the highest explanatory power (R^2^ 0.55, p-value = <0.0001), but the ability to form new aggregates was an important predictor for leaf decomposition in both Spearman and pgls analysis (Fig. 4). When phylogenetic dependencies between isolates were considered in the pgls models for wood decomposition, use of xylan (R^2^ 0.42, p-value 0.0001) and CCUA (R^2^ 0.28, p-value 0.015) still had the strongest explanatory power.

**Fig. 4.**
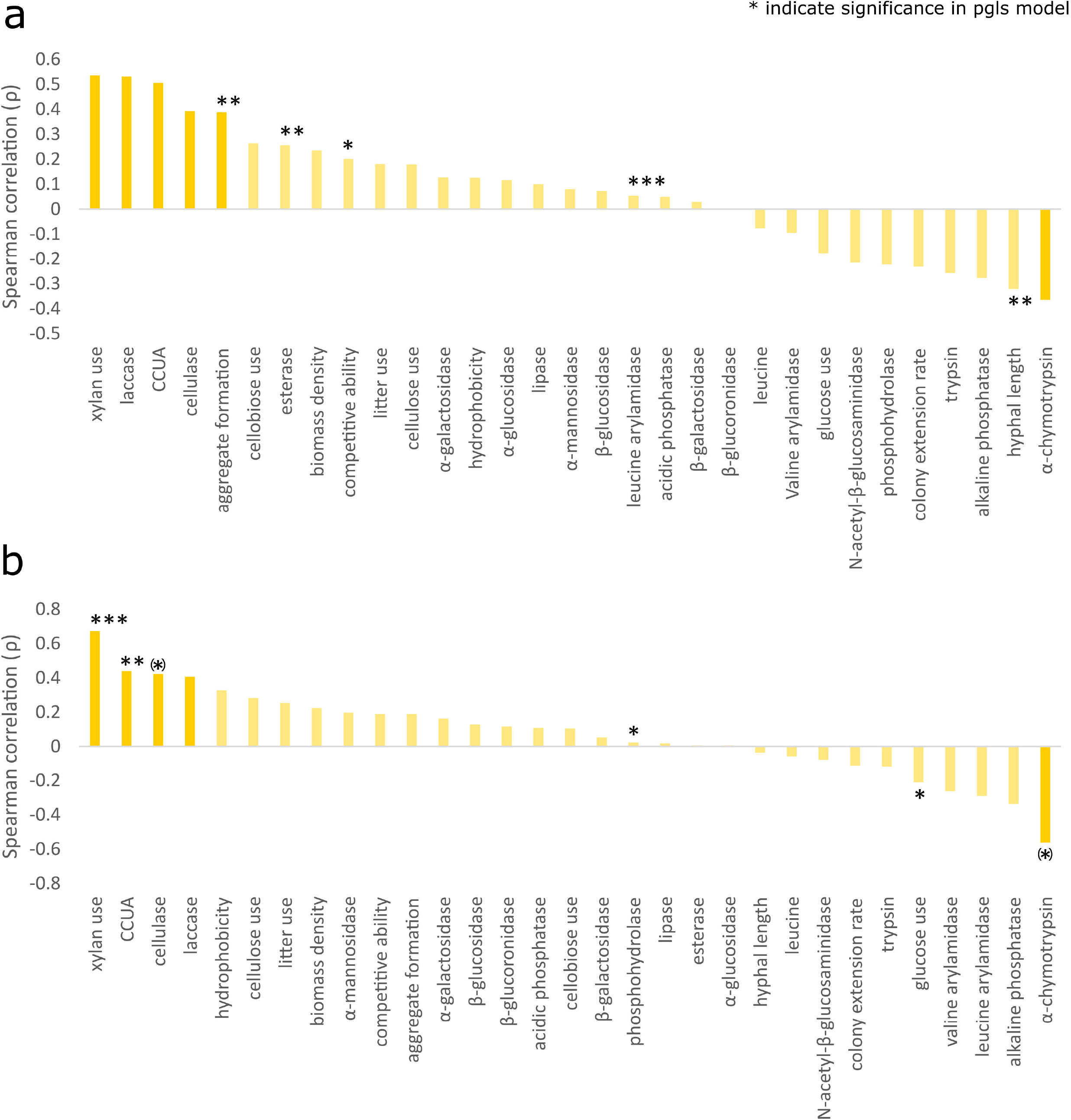

The PCA axis 1 explained 27% of the variance, while axis 2 explained 16 % of the variance (Fig. 5). The graph clearly showed that the Mucoromycota were completely separated from the other two phyla. Within the Ascomycota, average litter weight loss aligned well with cellulase activity and aligned with the ability to form new soil aggregates, the competitive ability, the ability to use xylan and more complex carbon sources (CCUA) as substrate. One isolate of the Basidiomycota (*Macrolepiota excoriata*, genus *Agaricales*) aligned with laccase activity and hyphal length. Activity of α-chymotrypsin, leucine arylamidase, phosphohydrolase and glucose use negatively aligned with litter weight loss.

**Fig. 5.**
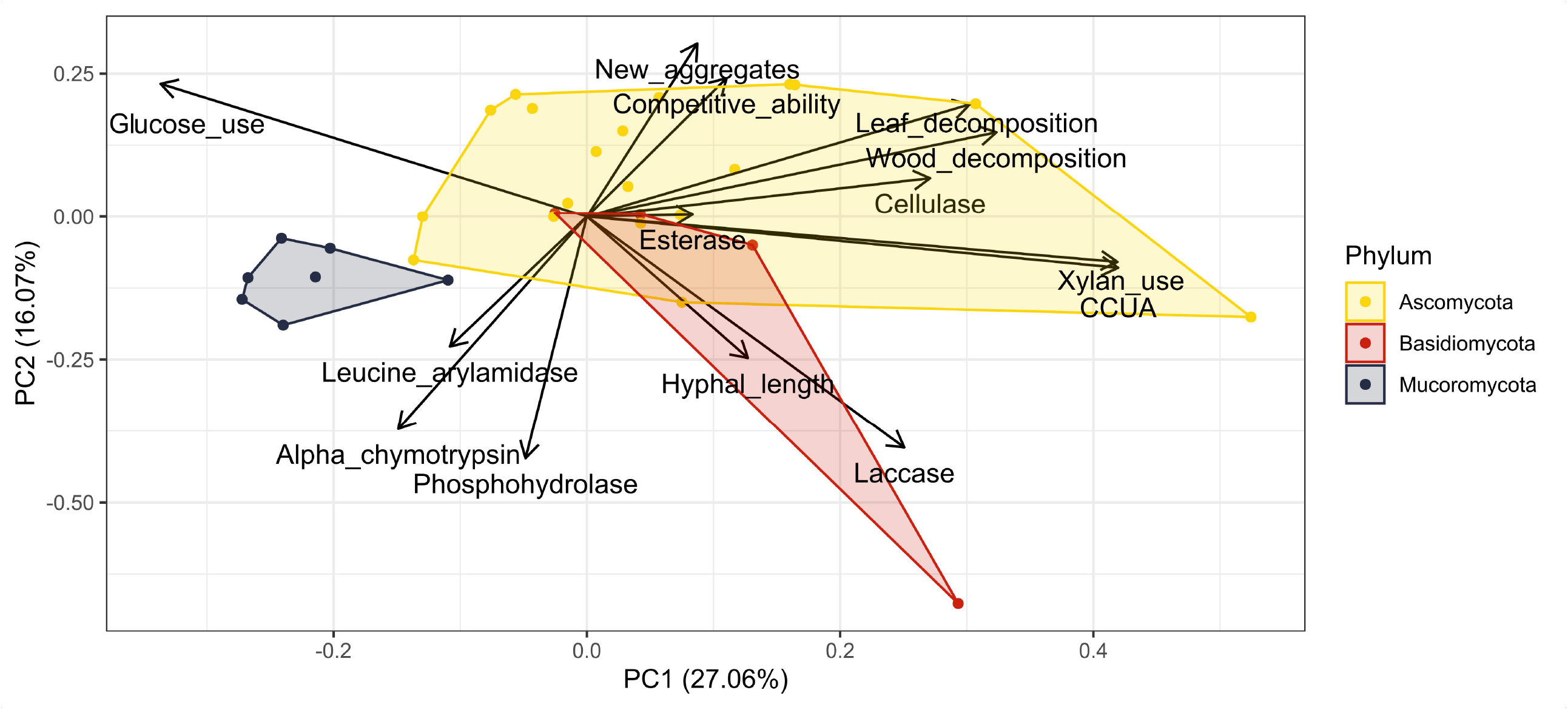

## Discussion

### Grassland soils harbor Ascomycota and Basidiomycota fungi with potential function in leaf litter decomposition

The functional ability to decompose litter varied substantially among isolates, with differences conserved at the phylum level. While within the Ascomycota and Basidiomycota most isolates showed the ability to decompose leaf litter, only 10 out of 24 isolates were also able to decompose wood. Mucoromycota had much lower ability to decompose leaf litter and showed no ability to decompose wood. The decomposition of wood litter was on average 41.8 % lower compared to leaf litter, showing that the complexity of litter is an important predictor for short-term decomposition rates in our grassland soil.

Leaves are generally considered ‘high quality’ litter that is decomposed fast while wood is considered ‘low quality’ litter that is usually decomposed more slowly (Cotrufo et al. 2013). This can be explained by litter chemistry, as leaves have more easy to decompose components (e.g. cellulose, hemicelluloses) and more nitrogen compared to wood, which contains more lignin (Brinkmann et al. 2002). The decomposition of the *Arrhenatherum* leaves was different between the phyla, for example Mucoromycota decomposed leaf litter only very slightly (increase of 3.2 % only compared to control), while Ascomycota and Basidiomycota increased decomposition by 26 and 29 %, respectively, in relation to the control plates. Furthermore, the fungi were isolated from a grassland soil and are probably mostly adapted to leaf litter from grassland plants, and they were grown in their home soil, thereby facilitating the decomposition of this litter type. However, grassland soils also harbor very complex organic matter, partly resulting from woody litter such as woody roots or woody stem pieces of certain shrubs and thus, wood litter also represents a valuable C source for soil fungi and adaptations are likely.

It has been shown before that certain substrates like cellulose can be preferred by a specific community where species are organized into functional guilds of decomposers (Bhatnagar et al. 2018). In addition to fungal preferences for certain substrates, some substrates require more complex processes for their degradation, e.g. the acquisition of mineral nutrients like N and P or other compounds such as proteins, fatty acids, or simple organic molecules (Algora Gallardo et al. 2021).

Interestingly, there is a connection between leaf decomposition and the ability to form new soil aggregates. As we showed in a previous study by Lehmann et al. (2020) the ability to form new aggregates is strongly influenced by fungal phylogeny, with Ascomycota being the best and Mucoromycota the worst aggregators. Based on their analysis of fungal trait contributions to soil aggregation the authors concluded that a critical local colony biomass density, rather than overall soil hyphal length, contributes to the formation of new soil aggregates. As mentioned above, the isolates of Mucoromycota are the weakest leaf decomposers in this study, while Asco- and Basidiomycota are effective decomposers. We also could not find a positive influence of hyphal length on decomposition. Instead, we also see a strong influence of the phylogeny of the data in leaf litter decomposition (but not in wood decomposition) and therefore conclude that phylogeny is a better predictor for leaf decomposition than overall hyphal length in soil. Therefore, we think that local dense biomass cannot only increase formation of new soil aggregates but also decomposition of high quality litter such as leaves, which were in a mesh bag, i.e. in a local spot.

### Complex carbon and xylan utilization by fungi are associated with litter decomposition

The ability to use xylan and more complex substrates in general (CCUA) were positively correlated with leaf and wood litter decomposition. Within our set of fungi we observed strong differences between the phyla but also large variations between isolates for this trait, showing that there are likely specialists among our set of fungi that are able to use one resource more effectively than others. This corroborates previous findings showing that members of the Mucoromycota, Ascomycota and Basidiomycota can have very different resource utilization capabilities (Hanson et al. 2008; Khosravi et al. 2015; Osono and Takeda 2002; Pawłowska et al. 2019).

Xylan is one of the main components of plant cell walls in grasses and wood. Thus, typically 20-30% of the grass cell wall contains this hemicellulosic polysaccharide (Hatfield et al. 2017), and in wood litter it can make up more than 30% of dry cell walls (Bajpai 2009). We found that the relation of xylan use and CCUA, respectively, with wood litter decomposition was conserved when we considered fungal phylogeny, showing that the correlation between wood decomposition and xylan use and CCUA is not only driven by fungal phylogeny of the data. Thus, we can use the ability to process complex carbon and to use hemicellulose like xylan as predictors for wood decomposition for our set of grassland fungi. For leaf litter the influence of the phylogeny on decomposition is not as strong as for wood litter, likely because leaves are a less complex substrate that is accessible through different strategies. Contrary to growth on xylan, biomass gain on glucose was negatively related to wood decomposition, showing that fungi preferring a simple sugar as substrate did not degrade the comparatively more complex leaf and wood litter very well. In our study, this can be attributed to the Mucoromycota, which had their highest biomass on glucose amended medium, were unable to decompose wood litter and only slightly decomposed leaf litter. A recent study showed that Mucoromycota also can use a wide variety of carbon sources (Pawłowska et al. 2019). However, this study also found large differences in the ability to use carbon sources and the study was focused on species of Mucorales and Umbelopsidales, whereas in our study we only have one isolate of each of those groups, but five isolates of Mortierellales. Hence, the selection of fungi and their inherent variable ability to decompose certain carbon resources leads to different results among studies.

### Enzyme activity correlated with litter decomposition is driven by fungal phylogeny

For selected enzymes we found a correlation with litter decomposition, with cellulase activity having the clearest positive relation. Cellulose and a few related polysaccharides are easy to decompose components of litter that make up 35-45 % of secondary cell walls in dry matter of grasses (Vogel 2008), whereas in wood cellulose can make up to 40-50% of dry mass, depending on wood types (Sikora et al. 2018). Given the abundance of cellulose in plant biomass, finding a positive correlation between cellulase activity and leaf litter decomposition is unsurprising. However, we found a consistent relationship between cellulase activity and wood decomposition, while for leaf decomposition the correlation is partly driven by fungal phylogeny.

At the individual isolate level laccase activity is an important explanatory trait for leaf and wood litter decomposition. However, a high laccase activity in our set of fungi did not always translate into a high decomposition (the isolate with the highest laccase activity had only a low decomposition rate). For both litter types this relationship is driven by the phylogeny, i.e. phylogenetic correction resulted in a non-significant relationship, probably because certain taxa within our dataset have similar traits that can drive such a correlation.

Some enzymes were only relevant in our correlation analyses when the data were phylogenetically corrected (esterase, leucine arylamidase, phosphohydrolase). Analyzing these 20 enzymes out of potentially many hundreds of enzymes in soils (Dick and Kandeler 2005; Senwo 2020), it becomes obvious that it is difficult to use single enzymes as predictors for litter decomposition. Usually, there is a diverse fungal community present in the soil, which has a large complex of enzymes that can degrade cellulose and other substrates sequentially, i.e. usually more than one enzyme is needed to degrade one compound (Allison 2012; Moore et al. 2020). Thus, by relating one single enzyme activity to the decomposition of litter, we are only looking at one puzzle piece with limited explanatory power.

The way the data were generated for the analyses likely added to the lack of correlation of enzymes and decomposition. To measure fungal enzymatic capabilities a separate experiment was necessary where we could access the pure fungal material on PDA. Whereas we measured litter decomposition in a more realistic experiment using water agar and soil. These two setups surely provide different amounts of resources, which influences the fungal metabolism. Camenzind et al. (2021) showed that enzyme activity may only be apparent in the absence of simple sugars, and was thus not detected in fungi cultured on PDA. Similarly, in another study testing wood decomposition by fungi cultivated on malt extract agar, no clear positive correlation between litter decomposition and enzymatic activity was observed; instead, only three out of nine tested enzymes showed a negative relation with decomposition rate (Lustenhouwer et al. 2020). We also observed a negative relation between an enzyme (α-chymotrypsin) and the decomposition of both types of litter. In our study this effect is mainly driven by Mucoromycota and one isolate of the Ascomycota, which are able to produce this enzyme and simultaneously show low decomposition.

### Phylogenetic effects on decomposition differ between leaf and wood litter

In our study we used three different phyla with the Mucoromycota clearly forming an ecologically separate group. They have different enzyme profiles and different abilities to process carbon and decompose litter, i.e. they were among the worst decomposers. This result is in agreement with the general notion in the literature of Mucoromycota being “sugar fungi”, and having less enzymatic capability for C degradation than Asco- and Basidiomycota (Moore et al. 2020; Pawłowska et al. 2019). However, this very general grouping has also been recently questioned by Pawlowska et al. (2019). The authors screened the capability of 52 mucoralean strains to use different C sources using the Biolog phenotypic microarray system, and found surprisingly diverse metabolic activity. However, similar to enzymatic profiles, Biolog data represent no direct evidence for decomposition potential of complex litter types. On the other hand, Pawloska et al. (2019) mainly included members of Mucorales and Umbellopsidales, while we also tested several Mortierellales. Indeed *Mucor* (RLCS01) and *Umbelopsis* (RLCS19) showed slightly different traits than *Mortierella*, but still clearly grouped within the Mucoromycota (Fig. 3). Additionally, some species of Mucoromycota are specifically known to be able to degrade hemicelluloses (Dix and Webster 1995). These different outcomes show that our results strongly depend on the species we include in our study.

The ecological separation of the Mucoromycota in our dataset clearly drives the phylogenetic signal that we find in leaf litter decomposition and the correlations with this variable. We therefore assume that the specific isolates among the Mucoromycota in our grassland soil will not have a strong effect on leaf decomposition but rather the other species that we isolated.

One Basidiomycete of our set is a member of the family *Polyporales* (RLCS09), a group known to be wood decomposers (Money 2016). This was the only Basidiomycete that decomposed wood in our experiment. The other three Basidiomycetes of our set are in the group Agaricales and did not decompose wood, but they decomposed leaf litter. This supports the general assumption that Agaricales are mainly decomposers of higher quality litter and not typical wood decay fungi (Floudas et al. 2020). Contrary to our study, the study by Lustenhouwer et al. (2020) found that wood decomposition was highly influenced by traits such as extension rate, combative ability and moisture. The authors, having used Basidiomycetes that were isolated from fruiting bodies on dead wood, suggest that extension rate could be an easy proxy for prediction of wood decay ability of fungi. We only had 4 isolates of Basidiomycota in our dataset, the majority was composed of two other phyla, and our fungi had a different origin (soil instead of wood) and were cultivated differently (PDA/soil vs malt extract agar). However, when looking at leaf decomposition, we also found that traits such as competitiveness, hyphal length and aggregate formation were important for decomposition. This shows that it is important to differentiate between forest and grassland ecosystems with their differences in litter and soil properties.

Ascomycota were the largest group in our data set (20 out of 31 fungi) and were among the best decomposers, especially for leaf litter. This corroborates existing knowledge on these fungi, which are known to be involved in, for example, straw decomposition (Sordariales and Hypocreales; Ma et al. 2013), wood decomposition (Xylariales; Helotiales; Cedeño–Sanchez et al. 2020; Richter and Glaeser 2015), some orders living on wood and litter (Pleosporales; Zhang et al. 2012) or they are known to live in the organic and litter layer of forest soils (Chaetothyriales, Heliotiales; Baldrian et al. 2012). We therefore want to underline the importance of this phylum for the prediction of litter decomposition in grassland soils.

## Conclusion and further research

Our study shows important differences in the decomposition of wood and leaf litter by saprotrophic grassland fungi. The direct assessment of fungal use of more complex C sources like xylan and cellulose (CCUA) appears to be a strong predictor of litter decomposition. This trait directly relates to a complete enzyme apparatus rather than individual enzymes. Fungi are known to possess large enzyme complexes for the degradation of complex C substrates like cellulose (Moore et al. 2020), with different species using different types of enzymes. Therefore, assays of individual enzymes may overlook the C degradation potential of certain fungal isolates. Thus, future studies should either use more direct assessments of fungal carbon use abilities or look at larger sets of enzymes or enzyme production by whole communities, in order to predict how a specific litter type will be degraded.

We found that the phylogeny of fungi in our dataset was very important for the prediction of litter decomposition. There are likely specialists in wood decomposition in our grassland soil that could be indicative for the degradation of complex substrates in grassland soils. In order to fully understand the role of the phylogeny for decomposition in grassland soils future research should include fungi from different grassland ecosystems, i.e. further species, and more importantly, this research should look at the role of fungal richness and diversity for litter decomposition. Another important factor for follow-up studies is litter diversity, i.e. the presence of more than one litter type at a time, which can also be decisive for the decomposition rate (Pei et al. 2017).

## Acknowledgements

EFL acknowledges funding from the Deutsche Forschungsgemeinschaft (LE 3859/1-1). TC acknowledges funding by the Deutsche Forschungsgemeinschaft (465123751, SPP2322 SoilSystems). We thank Johanna Kückes for help with experimental lab work.

## Data availability

The full data sheet has been deposited at https://github.com/Dr-Eva-F-Leifheit/fungal-traits-decomposition.

