## Supplementary Info for "Fungal traits help to understand the decomposition of simple and complex plant litter"

*FEMS Microbiology Ecology – Ecology of Soil Organisms*

The following Supporting Information is available for this article:

Table S1

Table S2

Methods details S3

**Table S1:** Information on fungal isolates used in the experiments. For each isolate we give information for phylum, class, order, family, taxon name and DSMZ accession number (Deutsche Sammlung von Mikroorganismen und Zellkulturen (German Collection of Microorganisms and Cell Cultures GmbH, DSMZ))

| **strain ID** | **DSMZ accession number** | **phylum** | **class** | **order** | **family** | **taxon name^1^** |
| --- | --- | --- | --- | --- | --- | --- |
| RLCS01 | DSM100293 | Mucoromycota | Mucoromycetes | Mucorales | Mucoraceae | *Mucor fragilis* |
| RLCS02 | DSM100407 | Mucoromycota | Mortierellomycetes | Mortierellales | Mortierellaceae | *Mortierella sp.* |
| RLCS03 | DSM100285 | Mucoromycota | Mortierellomycetes | Mortierellales | Mortierellaceae | *Mortierella alpina* |
| RLCS04 | DSM100322 | Mucoromycota | Mortierellomycetes | Mortierellales | Mortierellaceae | *Mortierella sp.* |
| RLCS05 | DSM100403 | Ascomycota | Sordariomycetes | Hypocreales | Nectriaceae | *Fusarium sp.* |
| RLCS06 | DSM100400 | Ascomycota | Sordariomycetes | Sordariales | Chaetomiaceae | *Chaetomium angustispirale* |
| RLCS07 | DSM100284 | Ascomycota | Sordariomycetes | Xylariales | Bartaliniaceae | *Truncatella angustata* |
| RLCS08 | DSM100325 | Ascomycota | Sordariomycetes | Hypocreales | Nectriaceae | *Fusarium sp.* |
| RLCS09 | DSM100406 | Basidiomycota | Agaricomycetes | Polyporales | Polyporaceae | *Trametes versicolor* |
| RLCS10 | DSM100286 | Ascomycota | Dothideomycetes | Pleosporales | Pleosporaceae | *Alternaria sp.* |
| RLCS11 | DSM100289 | Mucoromycota | Mortierellomycetes | Mortierellales | Mortierellaceae | *Mortierella alpina* |
| RLCS12 | DSM100405 | Ascomycota | Sordariomycetes | Sordariales | Chaetomiaceae | *Chaetomium angustispirale* |
| RLCS13 | DSM100290 | Ascomycota | Sordariomycetes | Hypocreales | Nectriaceae | *Fusarium sp.* |
| RLCS14 | DSM100404 | Ascomycota | Dothideomycetes | Pleosporales | Didymellaceae | *Nothophoma sp.* |
| RLCS15 | DSM100402 | Mucoromycota | Mortierellomycetes | Mortierellales | Mortierellaceae | *Mortierella sp.* |
| RLCS16 | DSM100408 | Basidiomycota | Agaricomycetes | Agaricales | Pleurotaceae | *Pleurotus sp.* |
| RLCS17 | DSM100324 | Basidiomycota | Agaricomycetes | Agaricales | Entolomataceae | *Clitopilus sp.* |
| RLCS18 | DSM100287 | Ascomycota | Sordariomycetes | Hypocreales | Nectriaceae | *Fusarium gibbosum* |
| RLCS19 | DSM100331 | Mucoromycota | Umbelopsidomycetes | Umbelopsidales | Umbelopsidaceae | *Umbelopsis isabellina* |
| RLCS20 | DSM100329 | Ascomycota | Sordariomycetes | Hypocreales | Ophiocordycipitaceae | *Purpureocillium lilacinum* |
| RLCS21 | DSM100327 | Ascomycota | Dothideomycetes | Pleosporales | Cucurbitariaceae | *Pyrenochaetopsis leptospora* |
| RLCS22 | DSM100401 | Ascomycota | Dothideomycetes | Pleosporales | Phaeosphaeriaceae | *Paraphoma chrysanthemicola* |
| RLCS23 | DSM101519 | Ascomycota | Sordariomycetes | Hypocreales | Stachybotryaceae | *Paramyrothecium sp.* |
| RLCS24 | DSM100410 | Ascomycota | Sordariomycetes | Hypocreales | Clavicipitaceae | *Metarhizium marquandii* |
| RLCS25 | DSM100292 | Ascomycota | Sordariomycetes | Hypocreales | Bionectriaceae | *Gliomastix sp.* |
| RLCS26 | DSM100330 | Ascomycota | Leotiomycetes | Helotiales | Helotiaceae | *Tetracladium apiense* |
| RLCS27 | DSM100326 | Ascomycota | Sordariomycetes | Sordariales | Chaetomiaceae | *Chaetomium subspirilliferum* |
| RLCS28 | DSM100323 | Ascomycota | Leotiomycetes | Helotiales | NA | *NA* |
| RLCS29 | DSM100288 | Basidiomycota | Agaricomycetes | Agaricales | Agaricaceae | *Macrolepiota excoriata* |
| RLCS30 | DSM100291 | Ascomycota | Eurotiomycetes | Chaetothyriales | Herpotrichiellaceae | *Exophiala sp.* |
| RLCS31 | DSM100328 | Ascomycota | Eurotiomycetes | Chaetothyriales | Cyphellophoraceae | *Cyphellophora sp.* |
| RLCS32 | DSM100409 | Ascomycota | Sordariomycetes | Hypocreales | Nectriaceae | *Fusarium oxysporum* |

1: best resolved tree annotation passing 80% threshold of bootstrap approach

**Table S2:** Information on the medium used for experiment II: In vitro experiments on carbon use

| **Element** | **Compound** | **Compound concentration** | **Comment** |
| --- | --- | --- | --- |
| C | glucose (C6H12O6) | 5 g L^-1^ | pH needs to be adjusted |
|  | cellobiose (C12H22O11) | 4.75 g L^-1^ |  |
|  | xylan (C5H8O4) | 4.4 g L^-1^ |  |
|  | cellulose (C12H20O10) | 4.5 g L^-1^ |  |
|  | litter | 4.42 g L^-1^ |  |
| N | NH_4_NO_3_ | 0.503 g L^-1^ |  |
| P | NaH_2_PO_4_ | 0.183 g L^-1^ |  |
| Mg, S | MgSO_4_ | 0.134 g L^-1^ | for more solid media double the addition |
| K | KCl | 0.023 g L^-1^ |  |
| Ca | CaCl_2_ | 0.065 g L^-1^ |  |
| Fe | NaFeEDTA | 0.02 g L^-1^ |  |
| vitamine | Thiamine HCl | 1 mg L^-1^ |  |
| vitamine | Biotin | 0.05 mg L^-1^ |  |
| Mo | Na_2_MoO_4_ | 0.05 mg L^-1^ |  |
| Cu | CuSO_4_ | 0.01 mg L^-1^ |  |
| Mn | MnSO_4_ | 0.05 mg L^-1^ |  |
| B | H_3_BO_3_ | 0.05 mg L^-1^ |  |
| Zn | ZnSO_4_ | 5 mg L^-1^ |  |
| gelling agent | Phytagel | 2 g L^-1^ | for more solid media add >2.5 g L^-1^ |

Supporting Information S3

**Detailed description of fungal growth medium with different C substrates**

A fungal growth medium was designed to specifically test for the differential effects of C sources on fungal growth (Table S2). To allow testing only for C effects, a complex medium composition was needed following principles described by Camenzind et al. (2020). The growth medium described therein allows to test for the limitation of individual elements, since all other elements are provided in sufficient amounts based on values given in the literature. For the experiment with different C sources, the medium here was modified in order to avoid osmotic stress by adding elements in high concentrations, especially when comparing glucose media with more complex C source which differentially affect the osmotic potential. Still, it was relevant to provide all elements in sufficient supply. To achieve this balance, element contents in fungal tissues were measured for all 31 fungal isolates (the same isolates used in this study): Fungi grown on media with high element supply and conditions of C:N=20 (medium details described in Camenzind *et al.* 2020) were freeze-dried, and C and N contents determined with an Elemental Analyzer (EuroEA, HekaTech, Germany). P, Mg, Ca, K and S contents were analysed after aqua regia digestion (1:4 HCl:HNO3) by ICP-OES analyses (Optima 2100 DV, Perkin Elmer, Germany). Based on these data, element concentrations X in the medium (calculated as C:X medium contents) were adjusted based on the average molar C:X ratio measured in fungal tissues (see Table S1). MgSO_4_ concentrations were doubled to ensure the formation of a stable gel matrix with Phytagel. Concentrations of micronutrients are still based on recommendations for fungal media given in the literature (Hoefnagels 2005). For media preparation P and C sources were autoclaved separately (Camenzind *et al.* 2020). C substrates were added in equal molar C amounts as described in the main manuscript. Litter C content was determined with an Elemental Analyzer (EuroEA, HekaTech, Germany).

Using more complex C sources, we encountered several methodological challenges in pre-experiments. Usually fungal growth media are prepared using agar, which is added in comparably high concentrations (20 g L^-1^; by comparison 5 g L^-1^ glucose used in our design). Since agar is in fact nothing else than a complex carbohydrate, some fungi can use it as a C source for growth. Testing the use of more complex C sources like cellulose or xylan cannot be proven due to the high parallel use of agar, which was observed by high growth rates on control plates only containing agar without any C source. To avoid this issue, we used a gelling agent that can be supplied to media in lower concentrations, and added phytagel at lowest concentration possible, i.e., 2 g L^-1^. Phytagel, as every gelling agent, also represents a potential C source. To solve this issue, control plates without added C substrates were used and biomass values on control plates subtracted from biomass gained on each C source.
Another complication with using complex C sources represents the reduced solubility in the medium, which results in agglomeration of C sources at the bottom of the petri dish. To avoid this bias, plates were filled by adding two layers. A first layer with 7.5 ml was added and solidified in petridishes Ø 6 cm. Thereafter, a second thin layer (4.5 ml) was added on top, to allow proximity of fungal hyphae to all C sources.
To extract fungal biomass efficiently, each medium was covered by a layer of 1 µm mesh (Polyamid, Sefar AG, Heiden, Siwtzerland). Following the principle of using cellophane as a separation of fungi and medium, mesh size and thickness of this mesh allows diffusions of all elements to the fungus. Cellophane proved to be problematic with more complex C source, since the higher production of enzymes degrades the cellophane layer quickly.

Camenzind, T, et al. (2020), 'Trait-based approaches reveal fungal adaptations to nutrient-limiting conditions', *Environmental Microbiology,* 22, 3548-60.

Hoefnagels, Mariëlle H. (2005), 'Biodiversity of Fungi: Inventory and Monitoring Methods', *BioScience,* 55 (3), 282-83.
